# Walking delays auditory processing in humans alongside a global shift in aperiodic neural activity

**DOI:** 10.64898/2026.11.04.764442

**Authors:** Xinyu Chen, Shitao Chen, Xin Zhang, Liyu Cao

## Abstract

Natural sensory processing unfolds during movement, yet human electrophysiology is usually measured under enforced stillness. Here, we used mobile EEG to test whether walking changes the timing of auditory processing. Thirty adults listened to lateralised pure tones while walking freely or standing. Walking led to a delay in auditory N1 latency (< 10 ms). Control analyses ruled out the possibility that the N1 delay was contaminated by foot strike related noise in EEG. Multivariate decoding further indicated a later emergence of sound location information during walking in the subset of participants with reliable decoding in both states. Walking also produced a scalp-wide decrease in the aperiodic exponent of neural responses. A larger global exponent decrease was associated with a larger N1 delay across participants. Together, these findings show that natural walking reorganises the temporal dynamics of human auditory processing and link early sensory slowing to a global change in ongoing cortical state.

## Introduction

Sensory processing is usually studied while the movements of participants are strictly constrained. Yet, sensory systems evolved to operate during continuous interaction with the environment. Work in mouse visual cortex established that locomotion can increase response gain and reshape spatial integration (Ayaz et al., 2013; Niell & Stryker, 2010), while dissociating locomotion from arousal revealed partially distinct contributions of these behavioural variables to neural responses (Vinck et al., 2015). In humans, mobile EEG (electroencephalogram) has also shown that walking leads to increased N1 amplitude in response to a visual stimulus compared to standing (Chen et al., 2022a, 2022b; Di Bello et al., 2026). Furthermore, alpha power in visual cortical areas was also reduced during walking (Cao et al., 2020; Cao & Händel, 2019; Ehinger et al., 2014; Scanlon et al., 2019), indicating an increasein neural excitability (Jensen & Mazaheri, 2010). Thus, movement is not simply a source of measurement noise. It is a biological state that has profound influence on sensory processing.

Auditory processing is also tightly coupled to movement. Motor related signals reach auditory cortex before and during movement, providing a circuit substrate for suppressing auditory responses to movement related noise (Schneider et al., 2014). Yet, locomotion does not uniformly suppress auditory responses. In mice, primary and higher order auditory fields show different mixtures of suppression and enhancement (Henschke et al., 2021), and movement related modulation is heterogeneous across neurons, sound classes, and movement types (Morandell et al., 2024). Auditory cortical populations can jointly encode external sounds and locomotion speed, indicating that movement related activity can carry behaviourally relevant information rather than merely adding noise (Vivaldo et al., 2023). Locomotion can also gate the transmission of auditory signals to association cortex and thereby alter multisensory decisions (Choi & Lee, 2025). Together, these findings suggest that auditory processing during locomotion reflects a reorganization of sensory coding across cortical circuits rather than a unitary gain change.

Human evidence remains comparatively sparse. Mobile EEG studies have shown that walking reallocates attentional resources, including reductions in target related P3 responses under naturalistic movement demands (Debener et al., 2012; Ladouce et al., 2019; Scanlon et al., 2019). A recent work demonstrated walking trajectory modulates auditory response gains. Responses to the input from the left ear increased before a left turn, and a similar pattern for the right ear (Chen et al., 2025). Most work, however, has focused on response magnitude, oscillatory power, or behavioural performance. Much less is known about whether walking changes the timing of auditory responses in the human brain. This distinction is important because the auditory N1 reflects an early, composite stage of cortical processing whose latency is sensitive to the timing of afferent and state dependent cortical operations (Naatanen & Picton, 1987). A locomotion related latency shift would therefore reveal a temporal reconfiguration of sensory processing that cannot be inferred from amplitude modulation alone.

One candidate mechanism underlying the influence of locomotion on auditory processing is a change in the ongoing cortical state. For example, auditory cortical responses vary with arousal and membrane-potential state, and very high arousal associated with walking can reduce the gain and reliability of sound-evoked responses relative to intermediate arousal (McGinley et al., 2015). At the macroscopic level, cortical state is also reflected in the aperiodic component of the electrophysiological power spectrum. Parameterising spectra into periodic and aperiodic components provides an estimate of the aperiodic exponent, which captures the steepness of the broadband 1/f-like spectrum (Donoghue et al., 2020). Biophysical modelling and invasive data have linked spectral slope to changes in synaptic excitation-inhibition balance (Gao et al., 2017), although pharmacological work showed that the exponent is not a unique or universally reliable proxy for excitation-inhibition ratio (Salvatore et al., 2024). We therefore treat the exponent here as an empirical marker of global cortical state rather than as a direct measure of excitability.

In the present study, we used 64-channel mobile EEG to measure auditory processing while participants listened passively to pure tones during natural overground walking and standing. We quantified N1 and P2 latency with jackknife-based fractional-area measures that separately captured the full component and its rising slope, simulated the contribution of foot strike related activity to test whether movement artefact could explain any latency shift, and used multivariate decoding to estimate when sound location information became available. Finally, we parameterised the aperiodic spectrum and asked whether locomotion related exponent changes covaried with auditory latency shifts across individuals. We hypothesised that walking would alter the temporal dynamics of auditory processing and that individual differences in this timing effect would be associated with a global locomotion related change in ongoing cortical state.

## Methods

### Participants

Thirty participants were recruited from Zhejiang University (14 women; mean age = 23.27 years, SD = 3.23 years). All participants reported normal hearing in both ears and provided written informed consent before participation. The study was conducted in accordance with the Declaration of Helsinki (2013) and was approved by the Ethics Committee of the Department of Psychology and Behavioural Sciences, Zhejiang University ([2022]047). Participants received monetary compensation of 50 RMB per hour after the experiment. No a priori power analysis was used to determine the sample size. With N = 30, a two-sided paired-samples t-test at α = 0.05 has 90% power to detect an effect of approximately dz = 0.61 for the critical within-participant comparison of ERP latency.

### Experimental design and data collection

The experiment used a 2 (movement state: walking vs. standing) × 2 (sound location: left vs. right) × 2 (sound frequency: 1000 vs. 1375 Hz) within-participants design. Within each movement state, 1000- and 1375-Hz tones were presented to either the left or right ear, yielding four auditory stimulus types. Participants completed six blocks: three walking and three standing. Half completed the blocks in the order W1-S1-W2-S2-W3-S3, and the other half in the order S1-W1-S2-W2-S3-W3 (W, walking; S, standing). Each block contained 400 stimuli, comprising 100 presentations of each stimulus type in random order. Thus, each participant received 2400 tones in total, with 1200 tones in each movement state. Each tone lasted 100 ms, including 5 ms linear rise and fall ramps, and was generated at a sampling rate of 44.1 kHz. Sound level was adjusted to a comfortable listening level. Stimulus onset asynchrony varied randomly between 1 and 1.5 s. Participants were instructed to listen to the sounds without making an overt behavioural response.

During walking blocks, participants walked along a corridor (36 metres long and 2.4 metres wide) at a comfortable, self-selected pace. At the end of the corridor, they turned and continued in the opposite direction until the block ended. Walking speed and cadence were not constrained. Plantar force was recorded during walking with force sensors placed under the heels inside the shoes. Force signals from both feet were sampled at 1000 Hz and used to identify the onset of foot strikes. During standing blocks, participants stood naturally with both feet on the ground and looked straight ahead. Force was not recorded during standing.

EEG was recorded with a 64-channel mobile system (Smarting PRO X, mBrainTrain LLC, Serbia) at 250 Hz. Electrodes were arranged according to the international 10-20 system. A common mode sense active electrode at FCz served as the online reference. The impedance of all electrodes was reduced to below 20 kΩ before recording.

The experiment was controlled in MATLAB (The MathWorks Inc, USA) using Psychtoolbox-3 (Kleiner et al., 2007). Auditory stimuli were delivered through DT 770 PRO headphones (Beyerdynamic, Germany) connected to a low-latency Sound Blaster Play3 HIFI sound card (Creative Technology Ltd., Singapore). Stimulus events, EEG, and force signals were synchronised and recorded using Lab Streaming Layer (Kothe et al., 2025). Each experimental block was stored as a separate data file, yielding six files per participant.

### EEG data preprocessing

EEG analyses were performed in MATLAB using FieldTrip (Oostenveld et al., 2011), EEGLAB (Delorme & Makeig, 2004), and in-house scripts. For each participant, data from all six blocks were first concatenated so that a single independent component analysis (ICA) could be applied across movement states. To estimate ICA unmixing weights, the concatenated data were high-pass filtered at 1 Hz and low-pass filtered at 30 Hz using zero-phase windowed sinc finite-impulse response filters with Kaiser windows. Bad channels were excluded if more than 1% of samples exceeded ±400 µV or contained flat line periods (seven participants had one bad channel, one participant had two, and one participant had three). The concatenated data were then cleaned with artefact subspace reconstruction using the first 120 s of standing data as the calibration segment and a cutoff of 7 (Mullen et al., 2015), rereferenced to the common average, and segmented into 1 s epochs. Epochs exceeding a 3 SD outlier criterion were removed before adaptive mixture ICA was run with one model and a maximum of 2000 iterations.

For ERP analyses, the original data were reprocessed with a 0.1 Hz high-pass and 30 Hz low-pass filter. No new ICA decomposition was estimated. Instead, the ICA unmixing weights obtained from the 1-30 Hz data were applied to these data, and signals were projected back to sensor space after excluding bad ICA components. Bad ICA components were rejected if any of the three following criteria were met. First, ICLabel (Pion-Tonachini et al., 2019) was used to flag components with a probability > 0.8 of reflecting eye or muscle activity. Second, a dipole fitting with the EEGLAB standard template boundary element head model was used to flag components with residual variance > 0.15. Third, spectra of component time series were estimated separately for walking and standing using Welch’s method. The walking to standing power ratio below 2 Hz was computed for each component. Among the 10 components with the largest ratios, components with a ratio > 1.05 were flagged to reduce the contamination of walking related low frequency noise. After component rejection, 17.53 (SD = 4.57) components remained on average.

A separate preprocessing stream was used for the aperiodic spectral analysis. It followed the ERP pipeline with two changes. First, the data were high-pass filtered at 0.1 Hz and low-pass filtered at 50 Hz. Second, ICA components were not rejected on the basis of walking to standing low frequency power differences. This avoided defining the spectral analysis by a condition dependent low frequency rejection criterion. An average of 18.63 (SD = 4.66) components remained for the subsequent analysis.

### Quantification of ERP latency and amplitude

ERPs were baseline corrected using the [-0.2 0] s pre-stimulus interval. Latencies were quantified with a leave-one-participant-out jackknife procedure (Kiesel et al., 2008). For each stimulus type in each movement state, a grand average ERP was calculated repeatedly while omitting one of the 30 participants, yielding 30 jackknife latency estimates based on 29 participants each. Individual ERP latency (Li) was recovered using the transformation described by Smulders (2010), Li = N·J̄ - (N - 1)·J(-i), where J̄ is the mean of the N leave- one-out latency estimates and J(-i) is the estimate obtained when participant i was omitted. Two latency measures were calculated. For component latency, the full component was scored with a 50% fractional area measure (Luck, 2014). With N1, the analysis included the portion of the N1 waveform below -0.5 µV. With P2, the portion above 0.2 µV was used. For rising-slope latency, the same fractional area procedure was restricted to the rising portion of the component. With N1, for example, the least negative mean N1 peak across stimulus types and movement states was used to define common 20% and 80% amplitude thresholds, and the waveform between the first crossings of those thresholds was entered into the fractional area calculation. Using common thresholds ensured that latency comparisons across stimulus types and movement states were not contaminated by condition specific peak amplitude. Before numerical latency extraction, ERP waveforms were upsampled to 5000 Hz with MATLAB’s resample function to provide a finer temporal grid. ERP amplitude was quantified as the mean voltage in a 20 ms window centred on the condition specific peak.

N1 analyses used the average waveform across Cz, C1, C2, FC1, and FC2. P2 analyses used the average across Cz and CPz. These electrode were chosen based on the topographies of group average responses (Fig. 1 b,d).

**Figure 1.**
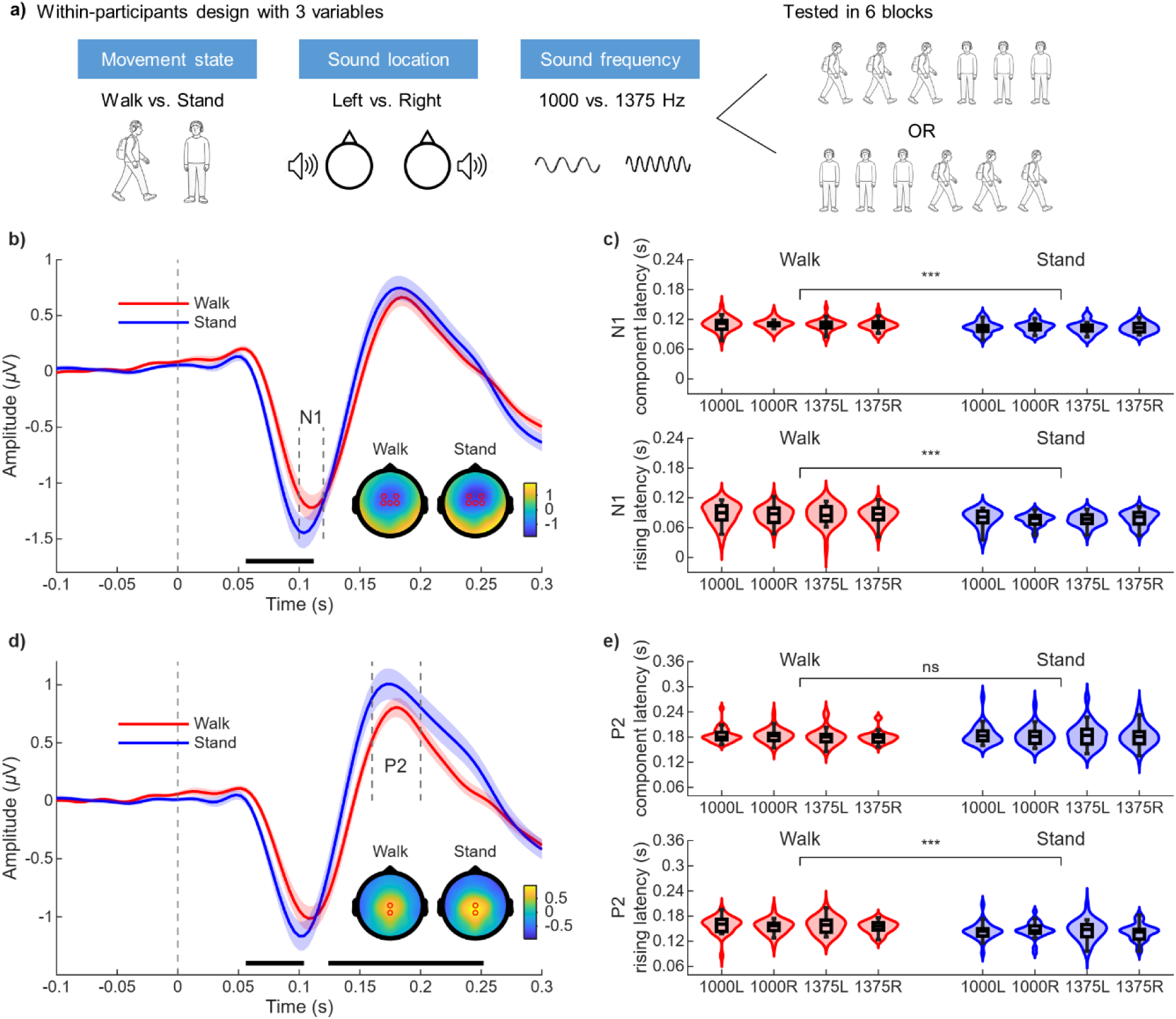
Walking delays auditory N1 response. (a) Experimental design. (b) N1 waveforms during walking and standing, averaged across the five highlighted electrodes. Horizontal bars indicate significant differences in amplitude between conditions. Topographies show the mean response from 100 to 120 ms. (c) N1 component latency (upper panel) and rising slope latency (lower panel) across experimental conditions. (d-e) Corresponding P2 waveforms and latency measures. *** *p* < 0.001, ns: not statistically significant.

### Strike ERP simulation analysis

Each foot strike produced a sharp increase in plantar force amplitude, which was used to detect strike onset. Strike onset locked ERPs were calculated separately for left and right foot strikes by averaging EEG data relative to strike onset. To estimate the contribution of strike ERP to the auditory ERP, we simulated a noise signal produced by strikes using the strike ERPs for every participant and electrode, using the actual temporal relationship between tones and foot strikes during walking. For each tone, the nearest four strikes were identified, with one left and one right strike before the tone, and one left and one right strike after the tone (Supplementary Figure 1a). The corresponding average left and right strike ERPs were shifted to their observed latencies relative to tone onset and summed to generate a single trial noise estimate (Supplementary Figure 1b). The single trial noise estimates were then averaged across trials using the same procedure as for the auditory ERP. The resulting average signal was the simulated noise signal, which was then added to the standing auditory ERP to create a noise simulated ERP. One participant was excluded from this simulation because of a technical error in the force data collection.

### Decoding analysis

Multivariate pattern analysis was used to quantify information about sound location (left vs. right) and sound frequency (1000 vs. 1375 Hz) in the EEG signal, separately for walking and standing. For sound location decoding, trials were pooled across sound frequencies within each location class. For frequency decoding, trials were pooled across locations within each frequency class. Single-trial EEG data were upsampled to 5000 Hz using the same procedure as in the ERP latency analysis, baseline corrected over [-0.2 0] s, and analysed from -0.1 to 0.3 s. For each movement state × stimulus dimension combination, cross validation used a leave-one-block-out scheme in which the three blocks of that movement state served as folds, ensuring that trials from the held-out block never entered the training set. Within each fold, trials from the same class were randomly grouped (group size = 10) and averaged into pseudo-trials to improve signal-to-noise ratio. Grouping was performed independently for the training trials and for the held-out test trials, with the number of pseudo-trials matched across classes. This random grouping was repeated 10 times, and the resulting decoding performance was averaged. Pseudo-trial features were z-scored using the mean and standard deviation of the training set only, which was then applied to the corresponding test set. A shrinkage regularised linear discriminant classifier implemented in MVPA-Light toolbox (Treder, 2020) was trained and tested independently at each time point. Decision values from each held-out fold were normalised using the mean and standard deviation of that fold’s own training-set decision values before being pooled across the three folds, and area under the receiver operating characteristic curve (AUC) was computed from the pooled decision values at each time point.

### Aperiodic spectral analysis

Power spectra were estimated separately for each experimental block. Preprocessed continuous EEG were divided into 4 s segments with 50% overlap, tapered with a Hamming window, and transformed to the frequency domain with a fast Fourier transform. Within each block, segments with extreme mean power from 1 to 30 Hz were removed using a median absolute deviation-median rule with a cutoff of 5 (Wilcox & Rousselet, 2023). Spectra from the remaining segments were averaged within block, and the three block spectra belonging to the same movement state were then averaged to obtain one walking and one standing spectrum per participant. Aperiodic activity was parameterised using FOOOF version 1.1.1 (Donoghue et al., 2020) via the MATLAB wrapper. Periodic peaks were modelled with peak-width limits of 0.5-15 Hz, a maximum of 6 peaks, a minimum peak-height of 0, and a peak threshold of 2. The remaining aperiodic component was fit in log-log space with the fixed model over 1-30 Hz. The resulting exponent quantifies the steepness of the 1/f-like spectrum, with larger values indicating a steeper slope. Model fit quality was assessed using the coefficient of determination (R^2^) and model fitting error. For each participant, the median R^2^ and fitting error were obtained across electrodes. No difference in R^2^ was found between walking (median R^2^ = 0.992; interquartile range: [0.982 0.996]) and standing (median R^2^ = 0.994; interquartile range: [0.989 0.996]; Wilcoxon signed-rank test, *W* = 158, *z* = -1.53, *p* = 0.125). However, smaller fitting errors (median error = 0.030; interquartile range: [0.023 0.035]) were found during walking than standing (median error = 0.038; interquartile range: [0.033 0.045]; Wilcoxon signed-rank test, *W* = 57, *z* = -3.61, *p* < 0.001).

### Statistical analyses

Time-resolved EEG amplitudes were compared across conditions using two-sided paired-samples t-tests or one-way repeated-measures ANOVA, as appropriate. For N1 and P2, data in the post-stimulus time window ([0 0.3] s) were tested, with multiple comparisons controlled using cluster-based permutation inference with 1000 Monte Carlo permutations (Maris & Oostenveld, 2007). ERP latencies were analysed with 2 (movement state: walking vs. standing) by 2 (sound frequency: 1000 vs. 1375 Hz) by 2 (sound location: left vs. right) repeated measures ANOVAs (Gladwin, 2020), with Greenhouse-Geisser correction when required. For group level decoding, AUC values across the whole decoding window were compared with the mean pre-stimulus AUC from -0.1 to 0 s using one-sided paired-samples t-tests, and false discovery rate was controlled at q = 0.01 (Benjamini & Hochberg, 1995; Benjamini & Yekutieli, 2001). For participant level decoding onset, sound location information was defined as available at the first post-stimulus time point that survived false discovery rate correction (q = 0.01) against a participant-specific null distribution generated from 1000 label permutations. For the aperiodic analysis, principal component analysis was applied to the participant × electrode matrix of walking-minus-standing exponent differences, and the first component score was used as the summary measure of the global exponent change. Bivariate outliers in correlation analyses were screened with the Robust Correlation Toolbox (Pernet et al., 2013). Since no outliers were identified, Pearson correlations are reported.

All statistical analyses were performed in MATLAB (The MathWorks Inc, USA). Unless otherwise specified, statistical significance was defined as *p* < 0.05.

## Results

### Walking delays early auditory processing

Participants listened to 1000 or 1375 Hz tones presented to the left or right ear while walking or standing (Fig. 1a). Auditory response timing was quantified with jackknife-based fractional-area measures applied either to the full ERP component or to its rising slope. For N1 (Fig. 1b,c), the three-way repeated measures ANOVA (movement state: walking vs. standing; sound frequency: 1000 vs. 1375 Hz; sound location: left vs. right) revealed a main effect of movement state comparing for both component latency (*F*(1,29) = 27.54, *p* < 0.001, *ηp*^2^ = 0.49) and rising slope latency (*F*(1,29) = 39.32, *p* < 0.001, *ηp*^2^ = 0.58). N1 was delayed during walking compared to standing by 6.20 ms (*SD* = 6.47 ms) for component latency and by 8.94 ms (*SD* = 7.81 ms) for rising slope latency. Sound frequency also affected N1 component latency (*F*(1,29) = 8.57, *p* = 0.007, *ηp*^2^ = 0.23), with earlier responses to 1375 Hz tones (*M* = 105.54 ms, *SD* = 10.25 ms) than to 1000 Hz tones (*M* = 107.40 ms, *SD* = 10.99 ms). No other N1 effects were significant. For P2 (Fig. 1d,e), movement state affected rising slope latency (*F*(1,29) = 42.38, *p* < 0.001, *ηp*^2^ = 0.59) but not component latency (*F*(1,29) = 0.36, *p* = 0.553, *ηp*^2^ = 0.01). P2 rising slope latency was delayed by 11.19 ms during walking (*SD* = 9.42 ms). No other P2 effects were significant. Walking was also associated with reduced ERP amplitudes (Supplementary Fig. 2a,b). N1 amplitude was smaller during walking (*M* = -1.19 µV, *SD* = 0.60 µV) than standing (*M* = -1.41 µV, *SD* = 0.74 µV; *t*(29) = 3.77, *p* < 0.001). P2 amplitude was likewise smaller during walking (*M* = 0.78 µV, *SD* = 0.44 µV) than standing (*M* = 0.99 µV, *SD* = 0.73 µV; *t*(29) = -2.49, *p* = 0.019).

### Strike related noise could not explain the auditory processing delay

Walking introduces movement-related activity into scalp EEG, raising the possibility that the apparent auditory delay reflects residual artefacts. Of particular concern is the ERP response time locked to foot strikes. Although the sound stimuli were randomly presented (i.e. not time locked to foot strikes), the potential influence of strike ERP was evaluated through simulation analysis. Specifically, we first obtained both the strike ERP to left and right foot strikes (e.g. Fig. 2a). For each tone in the walking block, the four nearest strike ERPs were summed with the observed timing of strikes relative to the tone preserved to create a single trial noise estimate. Single trial noise estimates were averaged to give a simulated noise signal, which was then added to the auditory ERP response in the standing condition to create a noise simulate ERP. This was done separately for N1 and P2 responses. For both N1 (Fig. 2a-d) and P2 (Fig. 2e-h), the noise simulated ERP did not differ in latency from the original standing ERP but retained the latency difference relative to the walking ERP (Supplementary Table 1). The same pattern was observed for ERP amplitude (Supplementary Fig. 2c,d). Thus, the modelled contribution of foot strike related activity was insufficient to account for the auditory latency delay observed during walking.

**Figure 2.**
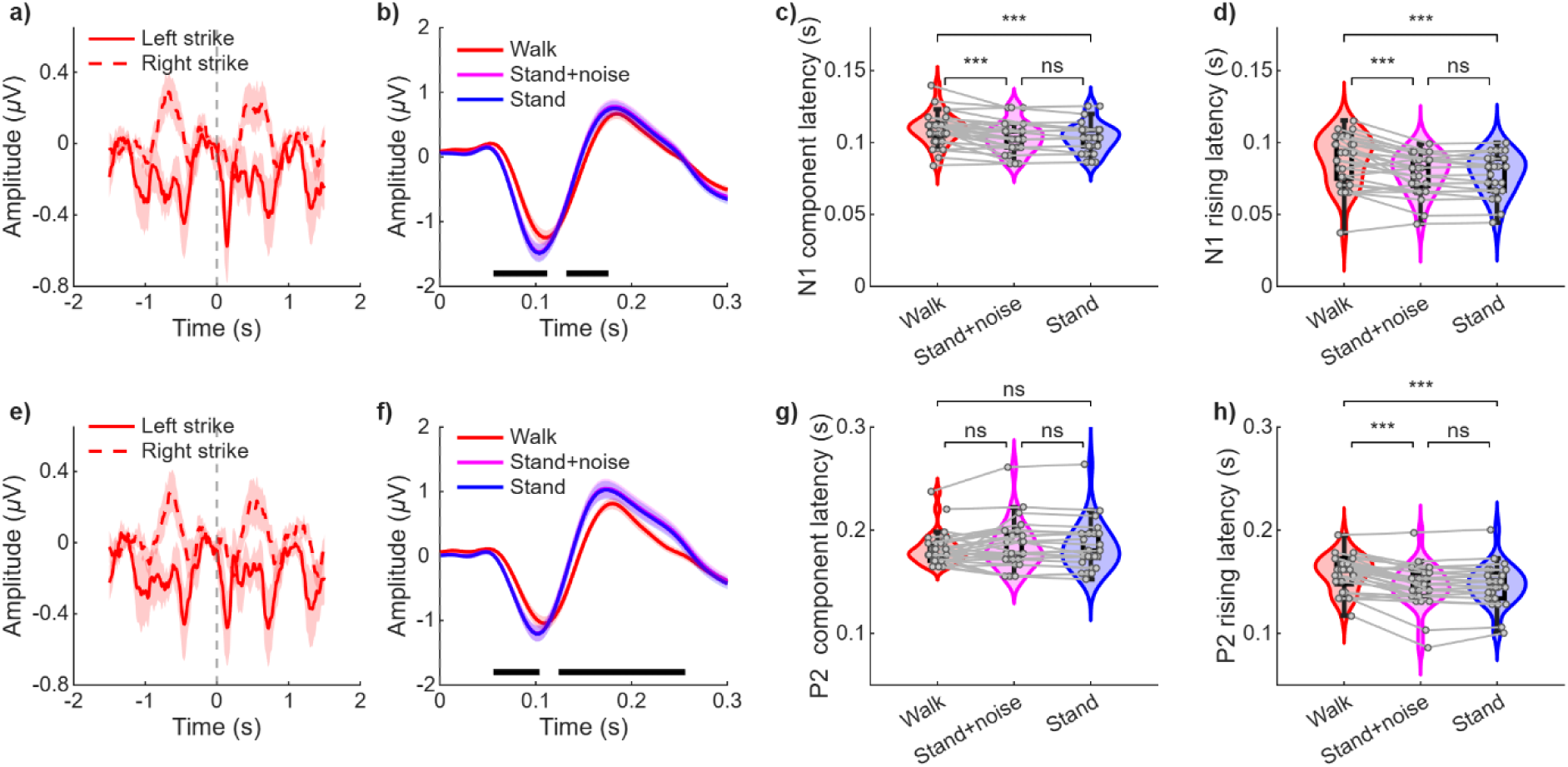
Foot strike related activity does not account for the auditory latency delay. (a) Foot strike ERP averaged across the N1 electrodes. (b) N1 waveforms during walking and standing, and the noise simulated waveform obtained by adding modelled strike related noise to the standing ERP. (c) N1 component latency. (d) N1 rising slope latency. (e-h) Corresponding analyses for P2, averaged across the P2 electrodes. *** *p* < 0.001, ns: not statistically significant.

### Walking delays the emergence of decodable sound location information

We next asked whether walking delayed the emergence of stimulus information in multivariate EEG patterns. Sound location (left vs. right) and sound frequency (1000 vs. 1375 Hz) were decoded separately in walking and standing conditions. The post-stimulus decoding results were compared to a baseline period ([-100 0] ms; before the sound onset) using paired-sample t-tests. Sound location was reliably decodable after stimulus onset (Fig. 3a), whereas sound frequency was not (Fig. 3b). For location decoding, significant post-stimulus time windows were found between [69 113] ms and [141 173] ms during walking and between [62 292] ms during standing. To compare information onset latency at the participant level, we identified the first post-stimulus time point with significant above chance location decoding. Reliable onset estimates in both movement states were available for 10 participants. Among them, the sound location information emerged later during walking (*M* = 83.79 ms, *SD* = 30.07 ms) than standing (*M* = 66.63 ms, *SD* = 13.16 ms; *t*(9) = 2.61, *p* = 0.029; Fig. 3c). This provides convergent evidence for delayed auditory information emergence during walking.

**Figure 3.**
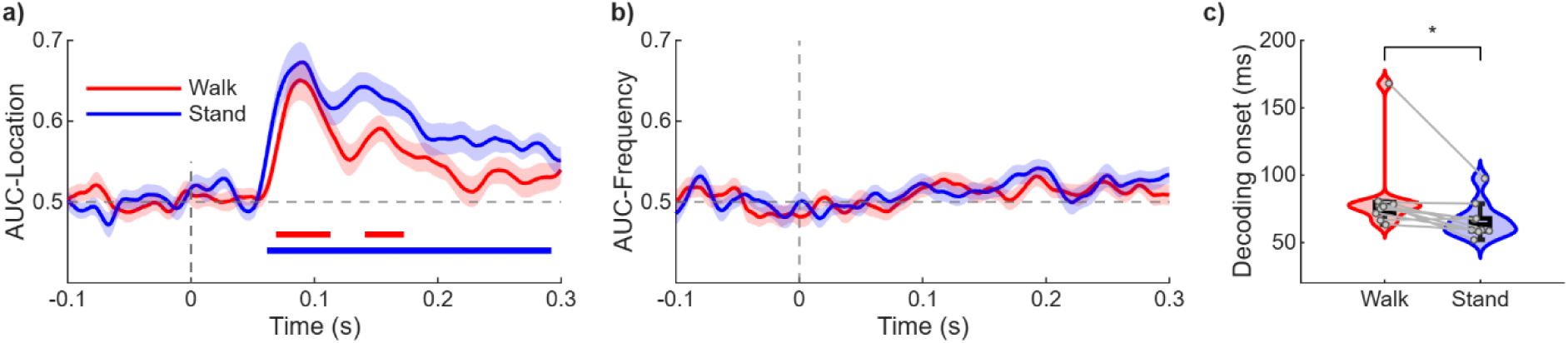
Walking delays the emergence of decodable sound location information. (a) Mean AUC for sound location decoding (left vs. right). Horizontal bars indicate post-stimulus intervals significantly above the pre-stimulus baseline. (b) Mean AUC for sound frequency decoding (1000 vs. 1375 Hz). (c) First post-stimulus time point with significant location decoding for participants with reliable decoding in both movement states.

### Correlation between slope changes of aperiodic neural activity and N1 delay during walking

The slope of aperiodic neural activity in each electrode was quantified using the exponent value, which was obtained via fitting the frequency domain EEG data after log transformation. The steeper the slope, the larger the exponent. A decrease in the aperiodic exponent was found for all electrodes during walking compared to standing (Fig. 4a). The exponent decrease during walking showed a global pattern across all the 64 electrodes, which was illustrated by the pairwise positive correlation of the exponent difference between walking and standing. Furthermore, a principal component analysis showed that the first component accounted for 76.5% of variance in the participant by electrode matrix of exponent difference (Fig. 4b). We therefore used the first component score to represent the global exponent change during walking. Interestingly, a significant negative correlation was found between the global exponent change and the N1 rising latency delay during walking (*r*(28) = -0.49, *p* = 0.006, Fig. 4c). This indicates that a larger decrease in aperiodic exponent during walking was associated with a larger delay in N1. No significant correlation was found between the exponent change and P2 latency change during walking (*r*(28) = -0.06, *p* = 0.736, Fig. 4d).

**Figure 4.**
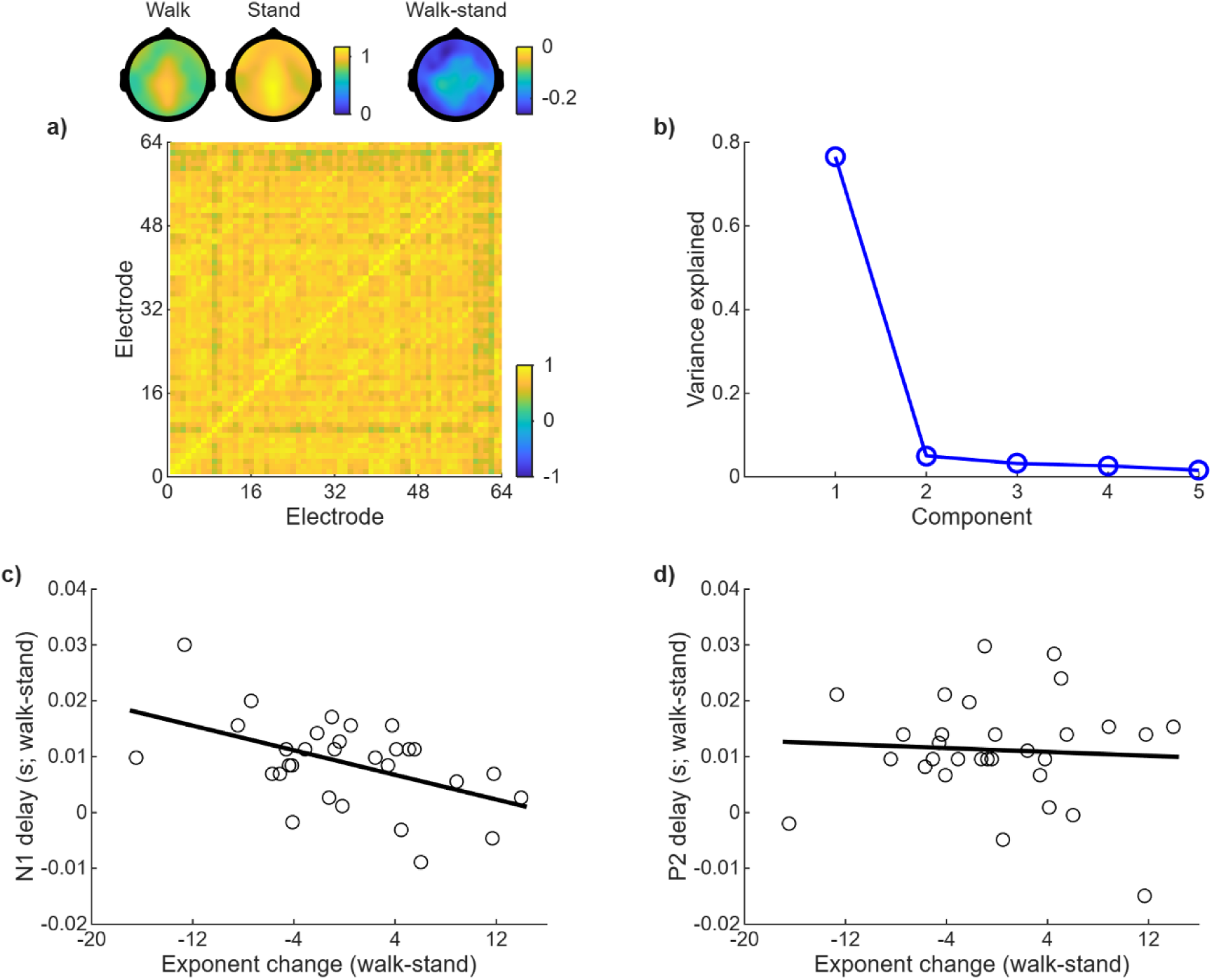
A global decrease in aperiodic exponent covaries with N1 delay. **(a)** Walking-minus-standing aperiodic exponent changes were positively correlated across the 64 electrodes. The topographies showed the exponent values during walking and standing, and the difference between the two. (b) Variance explained by the first five principal components of the participant by electrode exponent difference matrix. (c) Relationship between the global exponent change score and walking related N1 rising slope delay. (d) Similar to c, with P2 latency change during walking.

## Discussion

In the present study, we showed that compared to standing, walking led to a delay in auditory processing as indexed by the latency of N1 component in scalp EEG. This finding was corroborated by control analyses showing that foot strike related noise did not contaminate the processing latency estimation in the current experimental settings. Furthermore, results from the multivariate decoding showed that sound location information emerged later during walking. Walking also produced a scalp-wide decrease in the aperiodic exponent. Participants with larger global exponent decreases showed larger N1 delays. Together, these results suggest that locomotion reorganises the temporal progression of early auditory processing in humans and that this temporal reorganisation is embedded within a broader change in ongoing cortical state.

Previous work has established that movement can strongly alter the magnitude of auditory responses, although the direction of this modulation is mixed depending on the context. In mice, motor related signals can suppress auditory cortical excitatory neurons before and during movement through a motor-to-auditory corollary discharge pathway (Schneider et al., 2014). At the same time, locomotion related modulation is highly heterogeneous. Auditory responses can be either enhanced or suppressed across auditory cortical neurons and fields (Henschke et al., 2021; Morandell et al., 2024). Human mobile EEG studies likewise showed that walking changes auditory processing. Several auditory oddball studies have reported reduced P3 amplitudes during walking or increasingly demanding locomotor tasks, consistent with a redistribution of processing resources during movement (Debener et al., 2012; Ladouce et al., 2019; Reiser et al., 2019; Scanlon et al., 2019). Thus, locomotion does not impose a uniform gain change on the auditory system. The present findings add a distinct dimension to this literature. Irrespective of whether response amplitude is increased or reduced, the timing with which auditory information becomes available can itself be altered during locomotion.

The component specific pattern further suggests that walking did not simply translate the entire auditory ERP later in time. The N1 delay was robust for both the whole component and rising slope measures, whereas P2 showed a delay only for its rising slope and not for the latency of the full component. Auditory N1 reflects multiple, partly overlapping cortical processes associated with the early registration and analysis of acoustic events (Naatanen & Picton, 1987). The present scalp data cannot identify which generator or subprocess was delayed. Nevertheless, the convergence of two N1 latency measures makes an early cortical timing effect the most parsimonious interpretation. By contrast, the P2 rising slope is measured on the positive going transition that follows the N1 negativity and is therefore partly constrained by recovery from the preceding N1. A delayed N1 may consequently shift the apparent onset of the P2 rising phase even if the temporal centre of the P2 component is unchanged. The absence of a walking effect on full component P2 latency is potentially important. It raises the possibility that locomotion delays an early stage of auditory cortical processing without producing a uniform delay that accumulates through successive ERP components. In this sense, locomotion may selectively alter the temporal entry point of cortical auditory processing while later stages partially recover from the initial delay.

The direction of this auditory timing effect is particularly informative when considered alongside recent work in the visual system. Horrocks et al. (2024) showed that visual speed tuning in mouse V1 begins substantially earlier and persists for longer during locomotion, with population representations stabilising more rapidly than during stationary periods. This is almost the mirror image of the delayed auditory N1 and later sound location decoding observed here in humans. Such a difference should not be taken as evidence for a simple opposition between vision and audition, because both species and ecological demands differ. More generally, Gant and Mlynarski (2026) proposed that locomotion dependent visual modulation can be understood as an adaptive response to changes in the statistics of visual inputs. Their framework predicts different forms of modulation across receptive field scales, species and sensory environments. From this perspective, locomotion need not speed or slow all sensory systems in the same manner. Visual information required for online navigation may benefit from faster stabilisation, whereas auditory processing may be shaped by motor related gating, altered ongoing activity or the need to distinguish external sounds from predictable consequences of self-motion. A direct comparison of visual and auditory response latencies within the same freely walking human participants would therefore be an important next step.

The association between the aperiodic exponent and N1 latency provides a possible bridge between the behavioural state of locomotion and the timing of sensory processing. The exponent decrease was not restricted to auditory electrodes. The aperiodic exponent decrease was strongly correlated across the scalp, with the first principal component accounted for 76.5% of the spatial variance. The dominant effect therefore appears to reflect a spatially widespread change in electrophysiological state rather than a focal modulation, although further studies will be required to establish the anatomical extent of this change. Importantly, individual differences in this global exponent change were associated with the N1 delay but not the P2 latency change, linking the broadband state shift most closely to the early timing effect. Spectral parameterisation work has established the aperiodic exponent as a meaningful descriptor of broadband neural activity that varies with behavioural and cognitive state (Donoghue et al., 2020; for a recent review, see Hemmerich et al., 2026). Locomotion is a state when behaviour must remain responsive to continuously changing sensory and motor demands, rather than being optimised exclusively for the processing of a single sensory stream. That is, the cognitive system should be more flexible and variable during walking. Interestingly, recent work has linked the aperiodic exponent to metacontrol, that is, the dynamic regulation of processing between more persistent and more flexible cognitive states (Gao et al., 2024; Pi et al., 2024; Yan et al., 2024). They showed decreased aperiodic exponents in tasks requiring more cognitive flexibility (Yan et al., 2024) and increased aperiodic exponents in conditions requiring more persistent and selective processing (Gao et al., 2024; Zhang et al., 2023). Furthermore, Pi et al. (2024) showed that interindividual differences in resting state aperiodic activity predicted cognitive control styles, suggesting that aperiodic activity may index the brain’s capacity to adjust its processing mode to changing environmental demands. Our results fit very well with this line of reasoning that the scalp-wide exponent decrease observed during walking may reflect a transition towards a more flexible and variable cortical state. The slowing of early auditory processing may be a sign of such a state.

One caveat is that the present association between exponent decrease and N1 delay is correlational. It does not establish whether the aperiodic change contributes to the N1 delay, whether both are downstream consequences of locomotion related motor and arousal signals, or whether a third state variable drives both effects. A causal test will require experimental manipulation of the aperiodic exponent while measuring auditory timing. This possibility is becoming increasingly realistic because recent work has shown the possibility of targeting aperiodic EEG features directly with non-invasive stimulation (Masina et al., 2025; Stein et al., 2026).

In summary, natural walking alters when the human brain begins to process sound, not merely how strongly it responds. The robust N1 delay, the later emergence of sound location information, and the absence of a corresponding shift in full component P2 latency point to a temporally selective effect on early auditory cortical processing rather than a uniform slowing of the entire auditory cascade. At the same time, walking produced a spatially widespread flattening of the aperiodic spectrum. The magnitude of this aperiodic change tracked N1 slowing across individuals. Together with the broader literature on locomotion related sensory modulation, these findings support a view in which sensory systems are dynamically reconfigured during movement, with the consequences for response gain and timing depending on modality, processing stage, and behavioural demands. Establishing whether the aperiodic state plays a causal role in this reconfiguration is an important question for the next step.

## Data availability

The data are available upon reasonable requests to the corresponding author.

## Acknowledgments

This work was supported by National Natural Science Foundation of China (grant numbers: 32671392; 32271078).

## References

Ayaz, A., Saleem, A. B., Scholvinck, M. L., & Carandini, M. (2013). Locomotion Controls Spatial Integration in Mouse Visual Cortex. Current Biology, 23(10), 890–894. 10.1016/j.cub.2013.04.012

Benjamini, Y., & Hochberg, Y. (1995). Controlling the False Discovery Rate - a Practical and Powerful Approach to Multiple Testing. Journal of the Royal Statistical Society Series B-Statistical Methodology, 57(1), 289–300. 10.1111/j.2517-6161.1995.tb02031.x

Benjamini, Y., & Yekutieli, D. (2001). The control of the false discovery rate in multiple testing under dependency. Annals of Statistics, 29(4), 1165–1188. 10.1214/aos/1013699998

Cao, L., Chen, X. Y., & Haendel, B. F. (2020). Overground Walking Decreases Alpha Activity and Entrains Eye Movements in Humans. Frontiers in Human Neuroscience, 14. 10.3389/fnhum.2020.561755

Cao, L., & Händel, B. (2019). Walking enhances peripheral visual processing in humans. Plos Biology, 17(10). 10.1371/journal.pbio.3000511

Chen, X. Y., Cao, L., & Haendel, B. F. (2022a). Differential effects of walking across visual cortical processing stages. Cortex, 149, 16–28. 10.1016/j.cortex.2022.01.007

Chen, X. Y., Cao, L., & Haendel, B. F. (2022b). Human visual processing during walking: Dissociable pre- and post-stimulus influences. NeuroImage, 264. 10.1016/j.neuroimage.2022.119757

Chen, X. Y., Cao, L., Wieske, R. E., Prada, J., Gramann, K., & Haendel, B. (2025). Walking Modulates Active Auditory Sensing. Journal of Neuroscience, 45(45), e0489252025. 10.1523/JNEUROSCI.0489-25.2025

Choi, I., & Lee, S. H. (2025). Locomotion-dependent auditory gating to the parietal cortex guides multisensory decisions. Nature Communications, 16(1). 10.1038/s41467-025-57347-y

Debener, S., Minow, F., Emkes, R., Gandras, K., & De Vos, M. (2012). How about taking a low-cost, small, and wireless EEG for a walk? Psychophysiology, 49(11), 1617–1621. 10.1111/j.1469-8986.2012.01471.x

Delorme, A., & Makeig, S. (2004). EEGLAB: an open source toolbox for analysis of single-trial EEG dynamics including independent component analysis. Journal of Neuroscience Methods, 134(1), 9–21. 10.1016/j.jneumeth.2003.10.009

Di Bello, B., Ferrulli, N., Panacci, C., Filosa, M., Pitzalis, S., & Di Russo, F. (2026). The role of optic flow on reactive brain processing in cognitive tasks during locomotion. Brain Research Bulletin, 235. 10.1016/j.brainresbull.2026.111722

Donoghue, T., Haller, M., Peterson, E. J., Varma, P., Sebastian, P., Gao, R., Noto, T., Lara, A. H., Wallis, J. D., Knight, R. T., Shestyuk, A., & Voytek, B. (2020). Parameterizing neural power spectra into periodic and aperiodic components. Nature Neuroscience, 23(12), 1655–U1288. 10.1038/s41593-020-00744-x

Ehinger, B. V., Fischer, P., Gert, A. L., Kaufhold, L., Weber, F., Pipa, G., & König, P. (2014). Kinesthetic and vestibular information modulate alpha activity during spatial navigation: a mobile EEG study. Frontiers in Human Neuroscience, 8. 10.3389/fnhum.2014.00071

Gant, J. M., & Mlynarski, W. F. (2026). Locomotion optimizes sensory representations through a computational principle shared by rodents and primates. Science Advances, 12(35). 10.1126/sciadv.aed4172

Gao, R., Peterson, E. J., & Voytek, B. (2017). Inferring synaptic excitation/inhibition balance from field potentials. NeuroImage, 158, 70–78. 10.1016/j.neuroimage.2017.06.078

Gao, Y., Roessner, V., Stock, A. K., Mückschel, M., Colzato, L., Hommel, B., & Beste, C. (2024). Catecholaminergic Modulation of Metacontrol Is Reflected by Changes in Aperiodic EEG Activity. International Journal of Neuropsychopharmacology, 27(8). 10.1093/ijnp/pyae033

Gladwin, T. E. (2020). An implementation of N-way repeated measures ANOVA: Effect coding, automated unpacking of interactions, and randomization testing. Methodsx, 7. 10.1016/j.mex.2020.100947

Hemmerich, K., Santoni, A., Stottmeier, S., & Ronconi, L. (2026). Aperiodic EEG features of cognition: a systematic review. Reviews in the Neurosciences, 37(7), 689–726. 10.1515/revneuro-2026-0012

Henschke, J. U., Price, A. T., & Pakan, J. M. P. (2021). Enhanced modulation of cell-type specific neuronal responses in mouse dorsal auditory field during locomotion. Cell Calcium, 96, 102390. 10.1016/j.ceca.2021.102390

Horrocks, E. A. B., Rodrigues, F. R., & Saleem, A. B. (2024). Flexible neural population dynamics govern the speed and stability of sensory encoding in mouse visual cortex. Nature Communications, 15(1). 10.1038/s41467-024-50563-y

Jensen, O., & Mazaheri, A. (2010). Shaping functional architecture by oscillatory alpha activity: gating by inhibition. Frontiers in Human Neuroscience, 4. 10.3389/fnhum.2010.00186

Kiesel, A., Miller, J., Jolicæur, P., & Brisson, B. (2008). Measurement of ERP latency differences:: A comparison of single-participant and jackknife-based scoring methods. Psychophysiology, 45(2), 250–274. 10.1111/j.1469-8986.2007.00618.x

Kleiner, M., Brainard, D., & Pelli, D. (2007). What’s new in Psychtoolbox-3? Perception, 36, 14–14.

Kothe, C., Shirazi, S. Y., Stenner, T., Medine, D., Boulay, C., Grivich, M. I., Artoni, F., Mullen, T., Delorme, A., & Makeig, S. (2025). The lab streaming layer for synchronized multimodal recording. Imaging Neuroscience, 3. 10.1162/IMAG.a.136

Ladouce, S., Donaldson, D., Dudchenko, P. A., & Ietswaart, M. (2019). Mobile EEG identifies the reallocation of attention during real-world activity. Scientific Reports, 9. 10.1038/s41598-019-51996-y

Luck, S. J. (2014). An introduction to the event-related potential technique (Second edition. ed.). The MIT Press.

Maris, E., & Oostenveld, R. (2007). Nonparametric statistical testing of EEG- and MEG-data. Journal of Neuroscience Methods, 164(1), 177–190. 10.1016/j.jneumeth.2007.03.024

Masina, F., Napoli, E., Santacesaria, P., Giustiniani, A., Zago, S., Marino, M., Pezzetta, R., Montemurro, S., Manzo, N., Cona, G., Mapelli, D., & Arcara, G. (2025). Transcranial alternating current stimulation selectively modulates aperiodic EEG component: Unveiling alternative mechanisms of modulation. Clinical Neurophysiology, 177. 10.1016/j.clinph.2025.2110929

McGinley, M. J., David, S. V., & McCormick, D. A. (2015). Cortical Membrane Potential Signature of Optimal States for Sensory Signal Detection. Neuron, 87(1), 179–192. 10.1016/j.neuron.2015.05.038

Morandell, K., Yin, A. D., Del Rio, R. T., & Schneider, D. M. (2024). Movement-Related Modulation in Mouse Auditory Cortex Is Widespread Yet Locally Diverse. Journal of Neuroscience, 44(11). 10.1523/JNEUROSCI.1227-23.2024

Naatanen, R., & Picton, T. (1987). The N1 Wave of the Human Electric and Magnetic Response to Sound - a Review and an Analysis of the Component Structure. Psychophysiology, 24(4), 375–425. 10.1111/j.1469-8986.1987.tb00311.x

Niell, C. M., & Stryker, M. P. (2010). Modulation of Visual Responses by Behavioral State in Mouse Visual Cortex. Neuron, 65(4), 472–479. 10.1016/j.neuron.2010.01.033

Oostenveld, R., Fries, P., Maris, E., & Schoffelen, J. M. (2011). FieldTrip: Open Source Software for Advanced Analysis of MEG, EEG, and Invasive Electrophysiological Data. Computational Intelligence and Neuroscience, 2011. 10.1155/2011/156869

Pernet, C. R., Wilcox, R., & Rousselet, G. A. (2013). Robust correlation analyses: false positive and power validation using a new open source matlab toolbox. Frontiers in Psychology, 3. 10.3389/fpsyg.2012.00606

Pi, Y., Yan, J. M., Pscherer, C., Gao, S. D., Mückschel, M., Colzato, L., Hommel, B., & Beste, C. (2024). Interindividual aperiodic resting-state EEG activity predicts cognitive-control styles. Psychophysiology, 61(8). 10.1111/psyp.14576

Pion-Tonachini, L., Kreutz-Delgado, K., & Makeig, S. (2019). ICLabel: An automated electroencephalographic independent component classifier, dataset, and website. NeuroImage, 198, 181–197. 10.1016/j.neuroimage.2019.05.026

Reiser, J. E., Wascher, E., & Arnau, S. (2019). Recording mobile EEG in an outdoor environment reveals cognitive-motor interference dependent on movement complexity. Scientific Reports, 9. 10.1038/s41598-019-49503-4

Salvatore, S. V., Lambert, P. M., Benz, A., Rensing, N. R., Wong, M., Zorumski, C. F., & Mennerick, S. (2024). Periodic and aperiodic changes to cortical EEG in response to pharmacological manipulation. Journal of Neurophysiology, 131(3), 529–540. 10.1152/jn.00445.2023

Scanlon, J. E. M., Townsend, K. A., Cormier, D. L., Kuziek, J. W. P., & Mathewson, K. E. (2019). Taking off the training wheels: Measuring auditory P3 during outdoor cycling using an active wet EEG system. Brain Research, 1716, 50–61. 10.1016/j.brainres.2017.12.010

Schneider, D. M., Nelson, A., & Mooney, R. (2014). A synaptic and circuit basis for corollary discharge in the auditory cortex. Nature, 513(7517), 189–194. 10.1038/nature13724

Smulders, F. T. Y. (2010). Simplifying jackknifing of ERPs and getting more out of it: Retrieving estimates of participants’ latencies. Psychophysiology, 47(2), 387–392. 10.1111/j.1469-8986.2009.00934.x

Stein, A., Floegel, E., Zhang, M. S., Subramaniyan, A., Sharma, S., Feldman, Z., Evers, M., Riddle, J., Meltzer-Brody, S., & Frohlich, F. (2026). Establishing naturalistic brain stimulation targeting aperiodic EEG features: Transcranial endogenous current stimulation (tECS). Brain Stimulation, 19(2). 10.1016/j.brs.2026.103036

Treder, M. S. (2020). MVPA-Light: A Classification and Regression Toolbox for Multi-Dimensional Data. Frontiers in Neuroscience, 14. 10.3389/fnins.2020.00289

Vinck, M., Batista-Brito, R., Knoblich, U., & Cardin, J. A. (2015). Arousal and Locomotion Make Distinct Contributions to Cortical Activity Patterns and Visual Encoding. Neuron, 86(3), 740–754. 10.1016/j.neuron.2015.03.028

Vivaldo, C. A., Lee, J. Y., Shorkey, M., Keerthy, A., & Rothschild, G. (2023). Auditory cortex ensembles jointly encode sound and locomotion speed to support sound perception during movement. Plos Biology, 21(8). 10.1371/journal.pbio.3002277

Wilcox, R. R., & Rousselet, G. A. (2023). An Updated Guide to Robust Statistical Methods in Neuroscience. Current Protocols, 3(3). 10.1002/cpz1.719

Yan, J. M., Yu, S. J., Mückschel, M., Colzato, L., Hommel, B., & Beste, C. (2024). Aperiodic neural activity reflects metacontrol in task-switching. Scientific Reports, 14(1). 10.1038/s41598-024-74867-7

Zhang, C. Y., Stock, A. K., Mückschel, M., Hommel, B., & Beste, C. (2023). Aperiodic neural activity reflects metacontrol. Cerebral Cortex, 33(12), 7941–7951. 10.1093/cercor/bhad089

