## Supplementary materials for "Walking delays auditory processing in humans alongside a global shift in aperiodic neural activity"

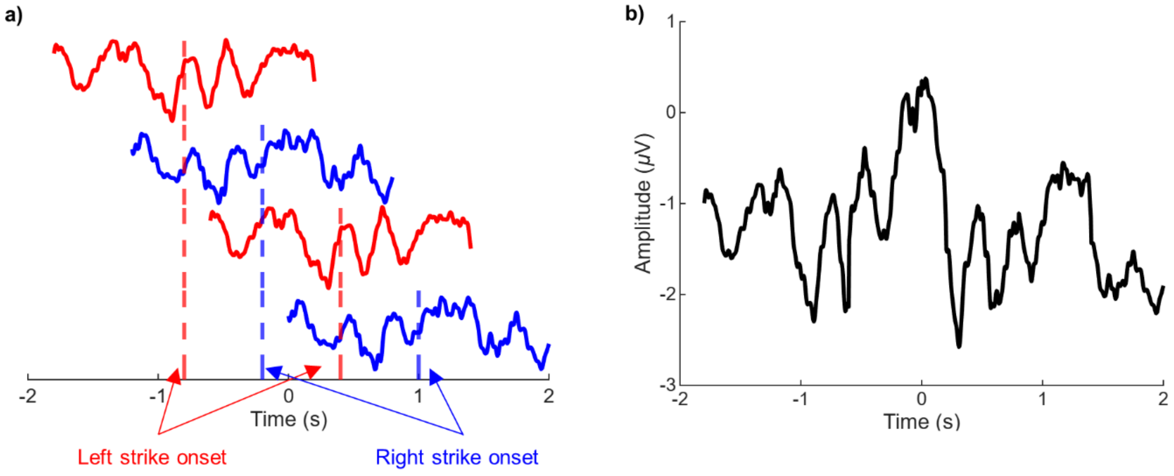


**Supplementary Figure 1. Strike related noise simulation.** (a) In this example, four strikes around a tone onset (time 0) were identified. The onset times of left strikes were -0.8 s and 0.4 s, and the onset times of right strikes were -0.2 s and 1.0 s. The red and blue lines represent the average foot strike related ERP from a particular electrode (Cz) of a single participant. (b) The single trial noise estimate obtained by adding the four waveforms in (a).

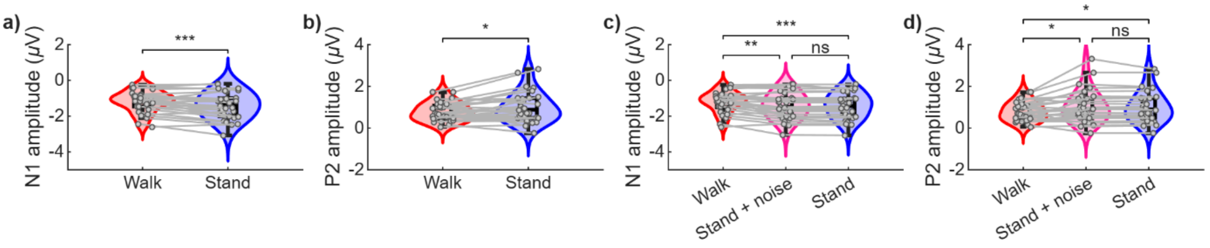


**Supplementary Figure 2. ERP amplitude comparisons.** (a) N1 amplitude was significantly lower during walking than during standing. (b) P2 amplitude was significantly lower during walking than during standing. (c) The noise simulated N1 showed no significant difference in amplitude with the standing condition ($t(28) = 1.61$, $p = 0.119$), but was significantly higher than the walking condition ($t(28) = -3.61$, $p = 0.001$). (d) The noise simulated P2 showed no significant difference in amplitude with the standing condition ($t(28) = 1.34$, $p = 0.192$), but was significantly higher than the walking condition ($t(28) = 2.56$, $p = 0.016$). N = 30 in a and b; N = 29 in c and d.

**Supplementary Table 1**

*Statistical comparisons of ERP latency in walking, standing, and noise-simulated data*

| | *Comparison* | *Mean Diff. (ms)* | *df* | *t* | *p* | *Cohen's d* |
|---|---|---|---|---|---|---|
| *N1 component latency* | | | | | | |
| | Walk vs. Stand + noise | 6.29 | 28 | 5.80 | < 0.001 | 1.08 |
| | Walk vs. Stand | 6.26 | 28 | 5.40 | < .0001 | 1.00 |
| | Stand + noise vs. Stand | -0.03 | 28 | -0.09 | 0.932 | -0.02 |
| *N1 rising slope latency* | | | | | | |
| | Walk vs. Stand + noise | 9.20 | 28 | 5.50 | < .001 | 1.02 |
| | Walk vs. Stand | 9.56 | 28 | 5.72 | < .001 | 1.06 |
| | Stand + noise vs. Stand | 0.36 | 28 | 0.70 | 0.489 | 0.13 |
| *P2 component latency* | | | | | | |
| | Walk vs. Stand + noise | -2.36 | 28 | -0.88 | 0.388 | -0.16 |
| | Walk vs. Stand | -1.68 | 28 | -0.63 | 0.537 | -0.12 |
| | Stand + noise vs. Stand | 0.68 | 28 | 0.96 | 0.344 | 0.18 |
| *P2 rising slope latency* | | | | | | |
| | Walk vs. Stand + noise | 12.60 | 28 | 5.84 | < .001 | 1.08 |
| | Walk vs. Stand | 12.11 | 28 | 5.85 | < .001 | 1.09 |
| | Stand + noise vs. Stand | -0.49 | 28 | -0.69 | 0.498 | -0.13 |
